# Personalized Regression Enables Sample-Specific Pan-Cancer Analysis

**DOI:** 10.1101/294496

**Authors:** Benjamin J. Lengerich, Bryon Aragam, Eric P. Xing

## Abstract

In many applications, inter-sample heterogeneity is crucial to understanding the complex biological processes under study. For example, in genomic analysis of cancers, each patient in a cohort may have a different driver mutation, making it difficult or impossible to identify causal mutations from an averaged view of the entire cohort. Unfortunately, many traditional methods for genomic analysis seek to estimate a single model which is shared by all samples in a population, ignoring this inter-sample heterogeneity entirely. In order to better understand patient heterogeneity, it is necessary to develop practical, personalized statistical models. To uncover this inter-sample heterogeneity, we propose a novel regularizer for achieving patient-specific personalized estimation. This regularizer operates by learning two latent distance metrics – one between personalized parameters and one between clinical covariates – and attempting to match the induced distances as closely as possible. Crucially, we do not assume these distance metrics are already known. Instead, we allow the data to dictate the structure of these latent distance metrics. Finally, we apply our method to learn patient-specific, interpretable models for a pan-cancer gene expression dataset containing samples from more than 30 distinct cancer types and find strong evidence of personalization effects between cancer types as well as between individuals. Our analysis uncovers sample-specific aberrations that are overlooked by population level methods, suggesting a promising new path for precision analysis of complex diseases such as cancer.

## 1: Introduction

A fundamental goal of pan-omic analysis, and a bottleneck for personalized medicine, is to understand the patterns of differentiation between individuals. With the advent of projects like The Cancer Genome Atlas^1^ (TCGA) and the International Cancer Genome Consortium (ICGC)^2^, genomic cancer data are generated at an unprecedented volume. We would like to use these data to understand patient-specific differences for personalized medicine, but many analysis pipelines discard sample heterogeneity in order to boost accuracy. Sample heterogeneity is particularly important for cancer, as cancer is increasingly appreciated as a complex disease in which many distinct underlying mutations may present with similar phenotypes (1); even within a single patient, there is increasing evidence of tumor mosaics composed of distinct cell lines (2). This difficulty with complex diseases like cancer motivates us to find new ways of analyzing data at increasingly small granularities.

Toward this aim, the bioinformatics community has developed increasingly specific assays (3). From targeted microarrays to whole-genome RNA-Seq and single cell RNA-Seq, the granularity of data collected by genomic assays has continued to be refined, to the point that we now possess data points representing the state of an individual cell at a single time point, unlocking the potential to study inter-patient, inter-tissue, and inter-cell variability of complex diseases.

A classic approach to personalization is to assume that we have access to a large volume of multimodal data (e.g. clinical, genomic, proteomic, biometric, etc.) on each individual, which is used to build large predictive models. Given enough data per individual, clinical outcomes and decisions can be personalized (3, 4), and recent work along these lines has leveraged a dizzying array of complex models including Gaussian processes (5), neural networks (6), and tree-based models (7), just to name a few. Despite the successes of these methods, they are still limited to this ‘one disease–one model’ perspective, in which a single predictive model—often through model averaging—is built for a single cohort (e.g. corresponding to patients of a particular disease type). Furthermore, these complex models are often difficult to interpret and are not guaranteed to provide correct inference into the underlying biological drivers of disease.

Unfortunately, in many circumstances, we may only have access to a limited amount of measurements per individual (e.g. either for cost or privacy reasons). In this case, it is advantageous to leverage data from distinct but related cohorts in order to build personalized models for each individual. For example, in cancer applications we now have access to large datasets for commonly studied cancers such as breast and lung cancer through repositories such as TCGA. At the same time, less common cancers such as of the eye and lymph node, have much less data (Table 1). A true “pan-cancer” study would combine all of this data, exploiting the similarities between different types of cancer to improve the accuracy of models for eye and lymph node cancer. That such similarities exist is well-established in the literature (e.g. 8). However, in the traditional ‘one disease–one model’ paradigm, data from other cancers play no role; while this makes sense for diseases which have a single root cause, the heterogeneity of complex diseases such as cancer renders these methods inadequate. Leveraging data from multiple cohorts while simultaneously obtaining distinct models for different diseases and different patients is a key challenge in personalized medicine.

**Table 1.**
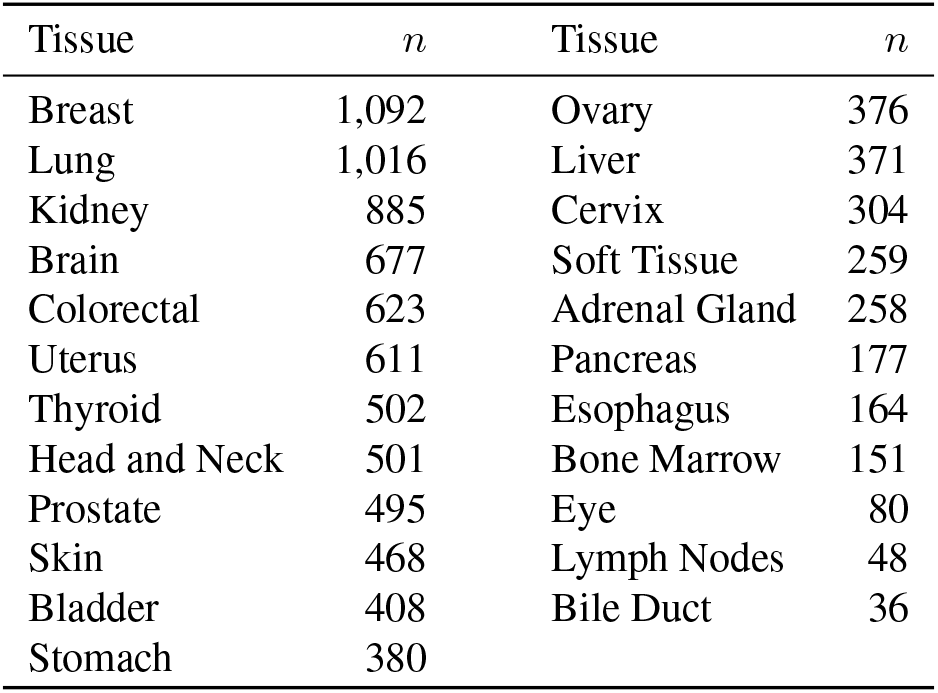
Number of samples by tissue in TCGA.

| Tissue | $n$ | Tissue | $n$ |
| --- | --- | --- | --- |
| Breast | 1,092 | Ovary | 376 |
| Lung | 1,016 | Liver | 371 |
| Kidney | 885 | Cervix | 304 |
| Brain | 677 | Soft Tissue | 259 |
| Colorectal | 623 | Adrenal Gland | 258 |
| Uterus | 611 | Pancreas | 177 |
| Thyroid | 502 | Esophagus | 164 |
| Head and Neck | 501 | Bone Marrow | 151 |
| Prostate | 495 | Eye | 80 |
| Skin | 468 | Lymph Nodes | 48 |
| Bladder | 408 | Bile Duct | 36 |
| Stomach | 380 |  |  |

Motivated by this new ‘many disease-many model’ paradigm, we propose a framework to estimate patient-specific models by learning patterns of differentation between samples. Instead of learning a single model for an entire cohort, our framework learns a unique model for each patient. The key is to leverage the fact that although each patient is expected to have a unique pattern of differentiation, these patterns are not independent of one another, and are expected to share substantial similarities. Leveraging this, we can “borrow strength” from the entire cohort to learn a useful model that is specific to a given patient. To do this, we propose a novel *distance matching* regularizer and show how it can be applied to regression problems. Our main contributions are three-fold:

1. A novel framework for personalized regression via distance matching;
2. We show that this framework can learn patient-specific models without prior knowledge of patient relatedness;
3. A TCGA pan-cancer study that illustrates the simultaneous similarities and differences between putative driver mutations of different cancers.

Although our main application will be to regression problems, our goal is *not* to simply predict an outcome, but instead to learn the underlying mechanisms that drive disease and lead to sample heterogeneity. By focusing our framework on learning patterns of differentiation, we can produce interpretable models of controllable granularity from patient-specific to pan-cancer.

## 2: Related Work

Traditional models assume only one or a few statistical parameters for a given population. As a simple example, consider the case of linear regression with response *Y* and predictors *X.* Then the typical model is *Y = × β + ϵ*, where the parameter *β* is shared across all individuals. More generally, mixture models allow for *K* different parameters *β*_*k*_, *k* = 1…,*K*. Another related generalization is multi-task learning (9). In both mixture modeling and multi-task learning, however, it is necessary that *K* ≪ *N* where *N* is the total number of samples in the cohort. By contrast, we are interested in the case *K* = *N*, i.e. a single parameterization for each individual in the cohort. This is what is meant by *personalized* regression models.

As it has long been a goal of biologists to understand inter-sample variation, significant prior work has aimed to estimate model parameters that vary between samples. Unfortunately, prior work requires either (1) a small number of sub-groups relative to the number of samples (e.g. mixture models) (10), or (2) known patterns of variation (11–13), or (3) significant domain knowledge to constrain the solutions (14, 15). Also closely related are random coefficient models, however, traditional random coefficient models do not allow for sample-specific sparsity patterns. In the presence of additional covariates *U* (often temporal or clinical variables), varying coefficient (VC) models have also been explored extensively (13, 16, 17). In this VC framework, each regression parameter is modeled as a function of some external covariates *U,* i.e. *β = f*(*U*). As with other models, VC models require significant domain knowledge in order to model a suitable relationship between *β* and *U*.

The closest work in spirit to ours is arguably the recent work on sample-specific network estimation (18, 19). Although these papers also consider the problem of sample-specific estimation, they focus on the particular problem of network estimation, and hence are not directly comparable to the present work.

## 3: Model

We are interested in learning which features 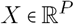 are relevant for predicting a phenotype 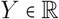 such as disease status. At the same time, we assume we have access to clinical covariates 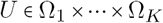 for each individual, which are allowed to be arbitrary—unordered or ordered, categorical or continuous, and even with missing values. Throughout, we let *N* denote the total number of patients in the cohort and use superscripts to identify samples. Thus, *Y*^(*i)*^, *X*^(*i)*^, and *U*^(*i*)^ denote the data for the *i*th sample and *β*^(*i*)^ denotes the personalized regression coefficients for the *i*th sample.

### A. Distance matching

To recover personalized model parameters *β*^(*i*)^ without *a priori* knowledge of how samples are related, we assume that there are *unknown* (pseudo)metrics *d*_*β*_ and *d*_*U*_ such that 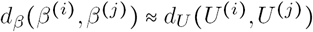. That is, similarity in parameters is related to similarity in covariates, however, the nature of this similarity is unobserved, unknown, and may not correspond to usual notions of distance such as Euclidean distance. This is closely related to the notion of distance metric learning introduced by Xing et al. (20). Existing work along these lines in the personalized estimation literature typically assumes that either (a) The metrics are Euclidean, or (b) The pairwise similarities are known (14, 15).

To learn these latent distance metrics, we model them as follows:

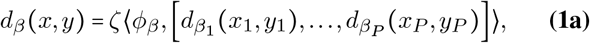

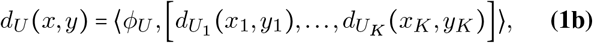

where ❬⋅,⋅❭ denotes the dot product of two vectors and *d*_*β*_ (*p* = 1,…,*P*) are user-specified metrics between scalars and 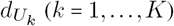 are user-specified metrics between covariates. Note that here we do not require these distance metrics to be differentiable. This allows for a wide variety of distance metrics, such as the discrete metric 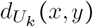 that equals one if *x = y* and is zero otherwise. This allows our framework to handle the realistic situation of categorical covariates without ordering. The parameters *ϕ**_ß_*** and *ϕ*_U_ represent unknown linear transformations of these “simple” distances into more useful latent distance metrics given by Eq. (1a) and Eq. (1b) with scale *ζ* > 0.

Define pairwise distance vectors for each *i, j* by

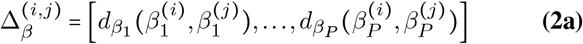

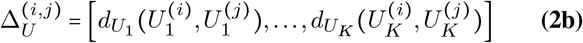

Since the covariate values in *U* are fixed, 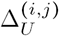 is also fixed, whereas 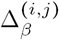 is not fixed since the values of *β*^(*i*)^ and *β*^(*j*)^ will change during training. For simplicity, we take 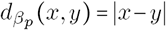 (*p* = 1,…, *P*) in the remainder of this paper.

Now define the following *distance matching regularizer:*

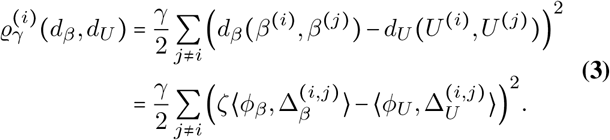

This regularizer attempts to match the pairwise distances between covariate values to the pairwise distances in the learned regression parameters. Let *f* be a loss function, e.g. least squares for regression or logistic loss for classification. Define a sample-specific objective by

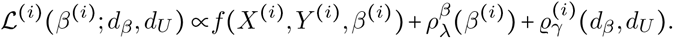

where *γ* trades off sensitivity to prediction of the response variable against sensitivity to sample distances, 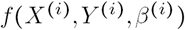 is the prediction loss for sample *i,* and 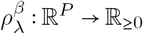 regularizes *β*^(*i*)^ with strength set by *λ*. Summing these, we obtain the complete objective function

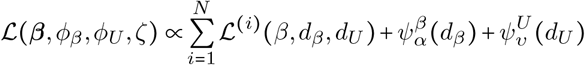

where 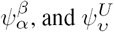 regularize the distance functions *d*_*β*_, *d*_*U*_ with strengths set by *α, υ,* respectively.

### B. Parametrization and initialization

Since the program Eq. (4) is nonconvex and the number of free parameters is large, some care must be taken to avoid degenerate solutions. We constrain the *ℓ*_1_ norm of both *ϕ*_*β*_ and *ϕ*_*U*_ to be equal to 1 and put all scaling into a single scale parameter *ζ*. In addition, we require that each entry of *ϕ*_*U*_ and *ϕ*_*β*_ is non-negative, ensuring non-negative distances between samples. Placing appropriate priors on *ϕ*_*β*_ and *ϕ*_*U*_, we arrive at the final program we wish to optimize:

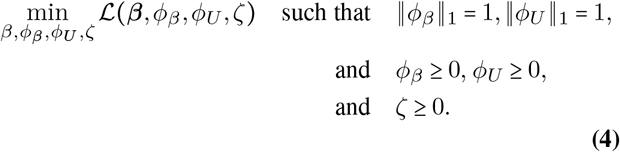

where inequality here is interpreted component-wise.

After normalization, the model Eq. (4) has (*N* + 1) *P + K* + 1 *free* parameters to be learned from *N* samples, which may seem significantly over-parameterized. Notwithstanding, although the technical details are beyond the scope of this short article, we can show that the distance matching regularizer Eq. (3) is able to constrain the personalized parameters 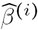 so that they do not deviate too far from a *population* regression estimation 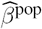, unless a substantial decrease in the loss can compensate for such deviations. Since Eq. (4) is a nonconvex program, proper initialization is crucial, and this gives us a practical strategy for initializing the personalized parameters: After solving for a population estimator 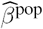, we initialize all 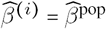. This initialization is important because the initial point is be a central point about which the personalized parameters are centered. As a result, our choice of regularizer allows for sample-specific personalization effects while preventing overfitting. This is a very desirable property for analysis of biological data: For example, suppose our data consists of microarray data from a diverse cohort of cancer patients. Each of these patients have experienced a series of mutations away from a healthy state; however, it is unlikely that they have experienced the same set of mutations. We would then like a personalized model to recover parameter values that are concentrated near a central model corresponding to a healthy state. This is precisely what distance matching does.

### C. Missing values

When there is a missing value in the covariate data, we set the distance between this value and all others to zero. This underestimates the distance between samples, biasing the solution toward retaining a central population estimator rather than personalizing the models based on missing features.

### D. Prediction

Although our main focus is on inference for a fixed sample cohort, given a new test point *X,* we can create a new model without re-running the learning algorithm on the entire dataset by averaging the personalized parameters of the *K* nearest models in the training set. This allows us to make predictions and inferences for new patients efficiently. Conveniently, since we have already learned a distance metric which we can use to accurately measure distance between samples, we can use this in the nearest neighbour search. Details are given in Algorithm 1. For linear regression, 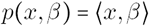. For logistic regression, 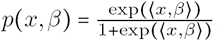.

## 4: Optimization

We seek to minimize Eq. (4) by first estimating a traditional regression estimator such as the Lasso or OLS, and then gradually relaxing the personalized regression models away from this population model. For simplicitly, we describe the procedure as centered about a single population estimator, however, the method trivially extends to initialization about a mixture model.

After setting hyperparameters *γ*, *α,* and *υ* (*λ* is dictated by the population estimator), we optimize by coordinate gradient descent with the following subgradients (note here that *Y*^(*i*)^ is a scalar value, and *X*^(*i*)^ and *β*^(*i*)^ *are* both *p*-vectors):

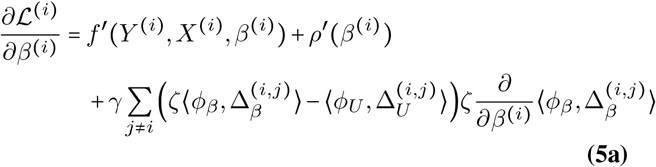

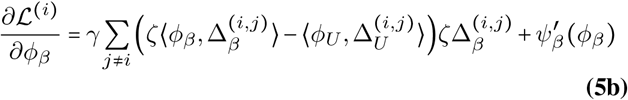

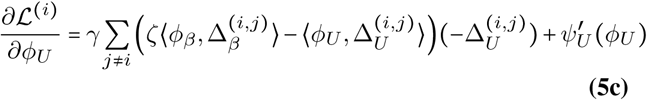

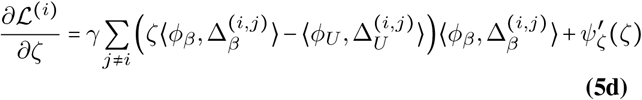

where 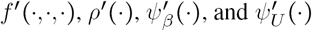 are subgradients of the predictive model *f*(·,·,·) and the regularizers 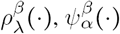, and 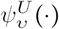, respectively.

The update to *β*^(*i)*^ is dependent on the distance metric chosen for parameter values. For 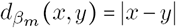, Eq. (5a) becomes

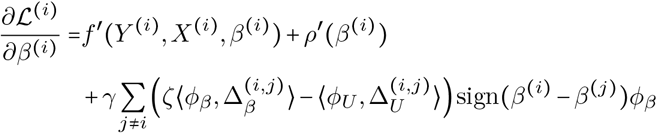

where the sign(·) function is applied element-wise. Finally, to ensure that each coordinate of *ϕ* is non-negative (i.e. distances cannot be negative), we project the updated value of *ϕ* into the non-negative reals. This is summarized in Algorithm 2. Each iteration of the naïve optimization procedure has computational time complexity in *O*(*N*^2^*PK*) where *P* is the number of features, *K* is the number of covariates, and *N* is the number of samples. This can be reduced to *O*(*NPK*) by defining a constant-size set of neighbors for each sample and only calculating pairwise distances within neighborhoods, as illustrated in Algorithm 2. The use of these neighborhoods naturally extends this procedure to optimization of personalized models centered around mixture models.

### A. Linear Regression

As an example application, let us instantiate the model Eq. (4) for personalized linear regression with Lasso regularization by

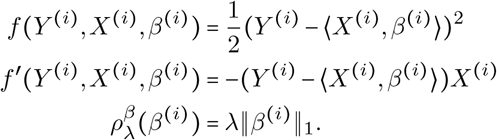

#### Algorithm 1 Inference Procedure

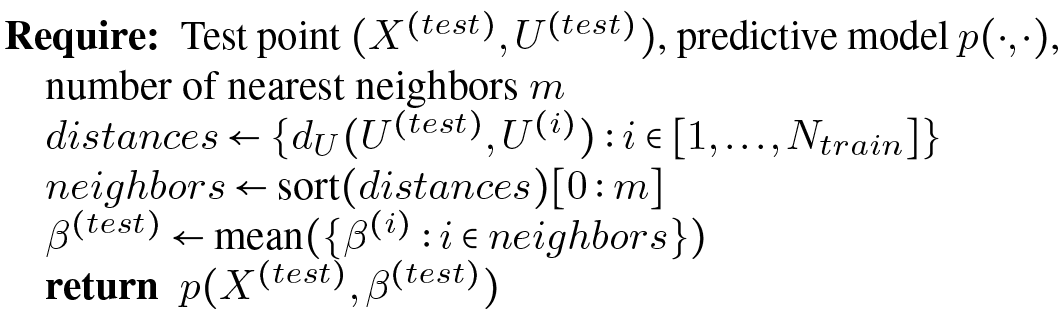

#### Algorithm 2 Optimization

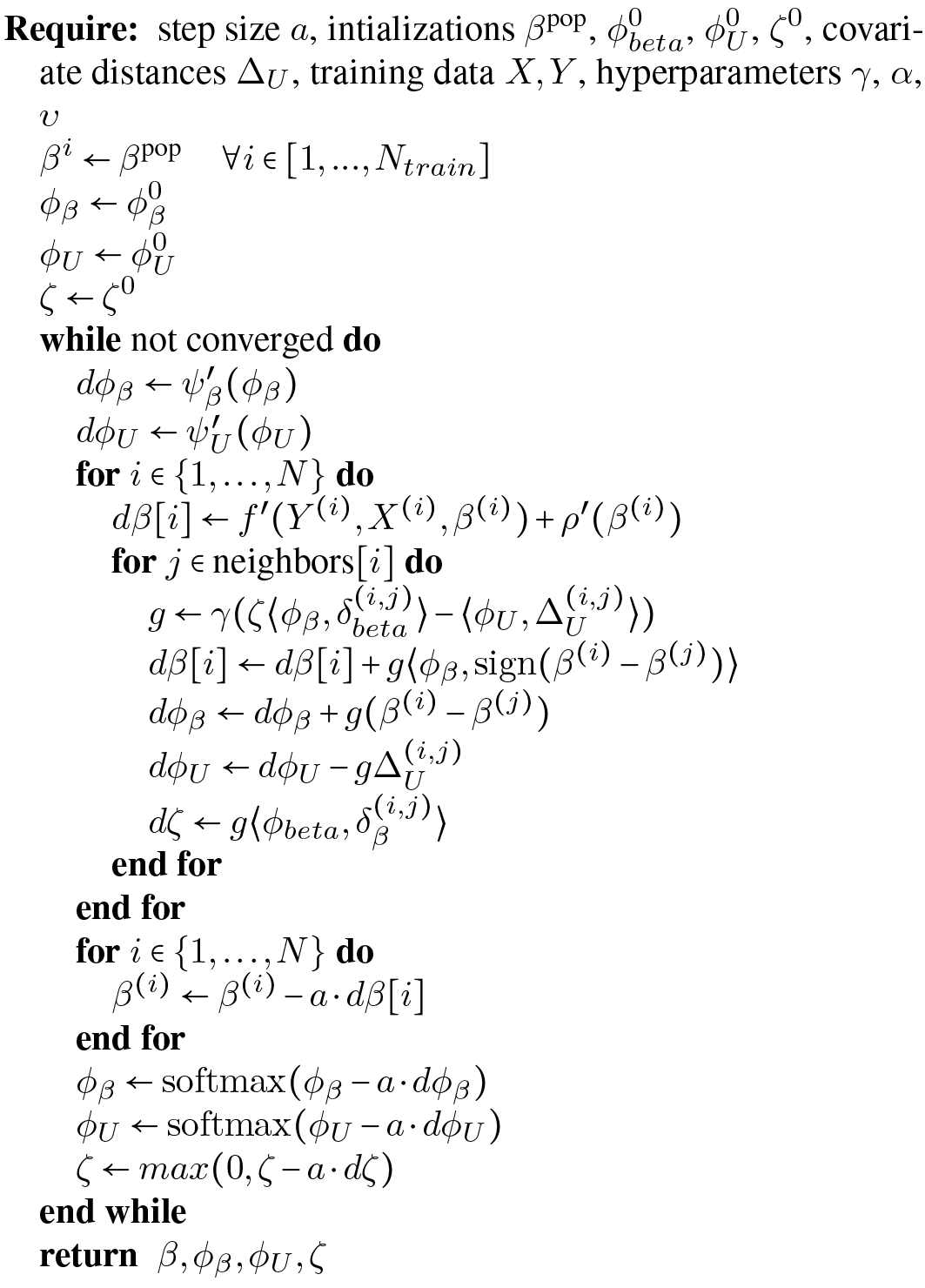

### B. Logistic Regression

Similarly, we instantiate personalized logistic regression with response variables 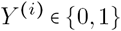 and Lasso regularization by

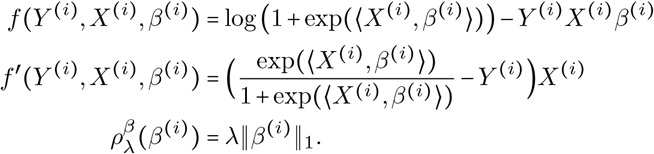

## 5: Simulation Study

To test the performance of personalized regression, we measure the recovery of personalized parameters on simulated data. For fixed 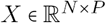, we generate sample-specific effect size vectors *β*^(*i*)^ ~ Unif (0,1) and sample *Y*^(*i*)^ ∈ {0,1} according to a logistic regression model. The covariates *U*^(*i*)^ are generated by projecting *ß*^(*i*)^ into *K < P* dimensions by multi-dimensional scaling. This produces covariates that are related to the personalized regression coefficients in a highly nonlinear, nonparametric manner. Recovery of the ground truth effect size vectors for fixed *P* = 10 variables and *K* = 3 covariates is depicted in Figure 1. In general, the personalized model outperforms other baselines except when the sample size is small, which is to be expected.

**Fig. 1.**
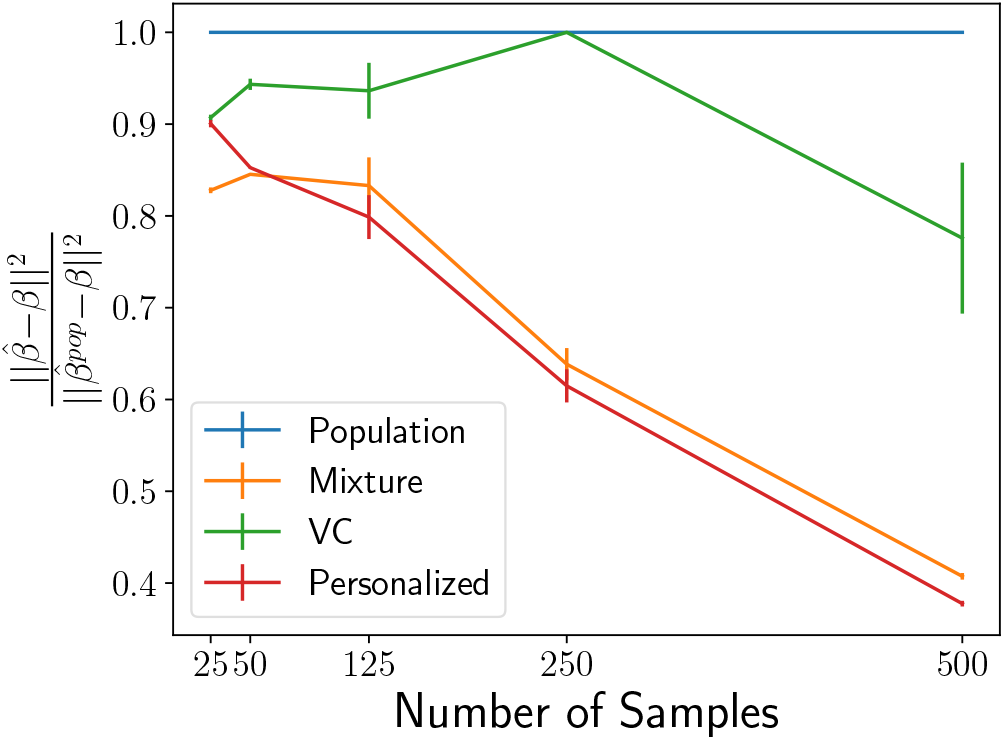
Recovery of regression parameters for the simulated data described in Section 5. Values indicate the mean error of the personalized parameter matrix normalized to the performance of the population estimator and averaged over 20 data generation processes, with error bars to denote the variance. The personalized model struggles at extremely low sample sizes but quickly surpasses the performance of the baseline models.

## 6: Sample-Specific Pan-Cancer Analysis

Here, we investigate the potential for personalized cancer analysis. We use gene expression (RNA-Seq) quantification data from The Cancer Genome Atlas (TCGA). This dataset compiles data from 37 projects spanning 36 disease types in 28 primary sites. After pruning for missing values, this dataset contains 9663 profiles for 8944 case and 719 matched control samples; we divide this set into 75% training data and 25% testing. While this full dataset is sizable, previous analyses have been hampered by the small number of samples for each particular cancer subtype (e.g., there are only 36 cases present in the bile duct cancer dataset). Because our framework of personalized regression allows models to share information across diverse settings, we are able to jointly analyze the cancer subtypes while still recovering subtype-specific characteristics. The number of samples available from each dataset was shown in Table 1.

We subsample genes based on annotations in the COSMIC Catalogue of Somatic Mutations in Cancer (21), so that there is exactly one putatively causal gene for each 5 non-annotated genes. This resulting in *P* = 4123 features when an intercept term is added. We train each logistic regression model to predict the case/control status of each sample with *ℓ*_1_ regularization to perform variable selection in order to study which genes are relevant for classification. Our baseline models include: *ℓ*_1_-regularized logistic regression model trained on all pan-cancer data (“Population”), *ℓ*_1_-regularized logistic regression model trained on each primary tissue type (“Tissue-Population”), *ℓ*_1_-regularized mixture model with the number of clusters equal to the number of tissue types in the pan-cancer dataset (“Mixture”), a logistic regression model with parameters that follow a linear varying coefficients model (“VC”), and the mixed model recently proposed by Hayeck et al. 22.

In addition to the RNA-seq data, we used the following 14 covariates: disease type, primary tumor site, age of the patient at diagnosis, year of birth of the patient, the number of days to sample collection, gender of the patient, race of the patient, percent of neutrophil infiltration, percent monocyte infiltration, percent normal cells, percent tumor nuclei, percent lymphocyte infiltration, percent stromal cells, and percent tumor cells in the sample. These covariates span a range of different types, including both continuous and discrete values; for continuous valued covariates, we use the *ℓ*_1_ distance function, for discrete valued covariates, we use the discrete distance metric. For the VC model, unordered discrete covariates such as primary tissue must be converted into one-hot vectors. This procedure increases the number of covariate features to 64, underscoring the benefit of our model’s ability to directly use the 14 unordered, discrete covariates without modification.

To predict case/control status of each sample, we implemented the personalized logistic regression model with Lasso regularization described in Section B. We selected *λ* in the population estimator by 10-fold cross-validation on the training set. This value of *λ* is held fixed between the population estimator and the personalized estimator. Next, we set *γ* so that the loss due to the distance matching regularizer is similar in magnitude to the prediction loss. Finally, we set *υ* and *α* so that the loss due to distance metric regularization is one order of magnitude smaller than the logistic classification loss. This heuristic represents our uncertainty in the form of personalization for cancer; we prefer to rely on the data than to set a rigid form of personalization. Empricially, we observe robustness in the solutions up to an order of magnitude change in these hyperparameters. By inspecting the variables (mRNA transcripts) selected by this method, we find that personalized regression identifies (1) individualized genetic aberrations, (2) interpretable patterns of differentiation, and (3) patient sub-typing that is more meaningful than clustering based on covariate data.

### A. Predictive Accuracy

To verify accuracy of the model, we first examine the classification loss of the case/control status target. Although our main goal is to study the selection of important genes for this task, the overall classification error is a convenient benchmark for sanity checking the learned models. Training and testing error values are shown in Table 2 with testing error rates calculated using *m* = 3 neighbors, as described in Algorithm 1. For the Tissue-Population model, we report the sample-weighted mean performance of the tissue-specific models. We see that the predictive accuracy for both training and testing sets is meaningfully improved by this method of personalization. We expect a low training error by virtue of the large number of parameters in the personalized models; the low testing error indicates that personalized patterns of differentiation are generalizable throughout the patient cohort and that the learned distant metrics are effective at finding related samples at test-time.

**Table 2.**
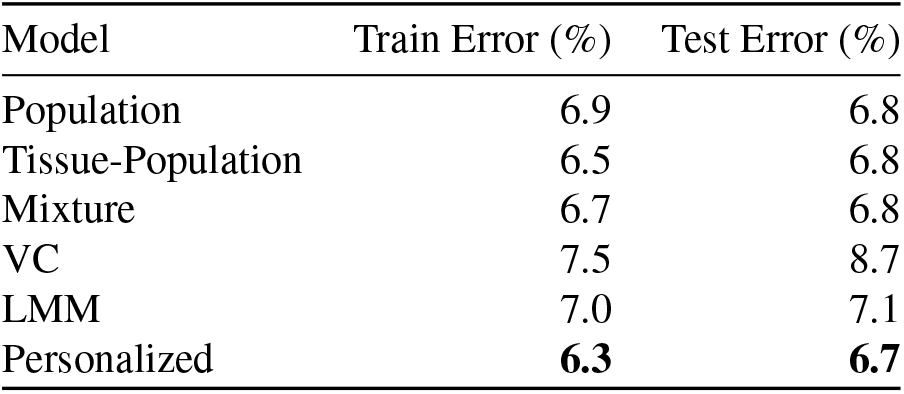
Classification Errors

| Model | Train Error (%) | Test Error (%) |
| --- | --- | --- |
| Population | 6.9 | 6.8 |
| Tissue-Population | 6.5 | 6.8 |
| Mixture | 6.7 | 6.8 |
| VC | 7.5 | 8.7 |
| LMM | 7.0 | 7.1 |
| Personalized | **6.3** | **6.7** |

**Fig. 2.**
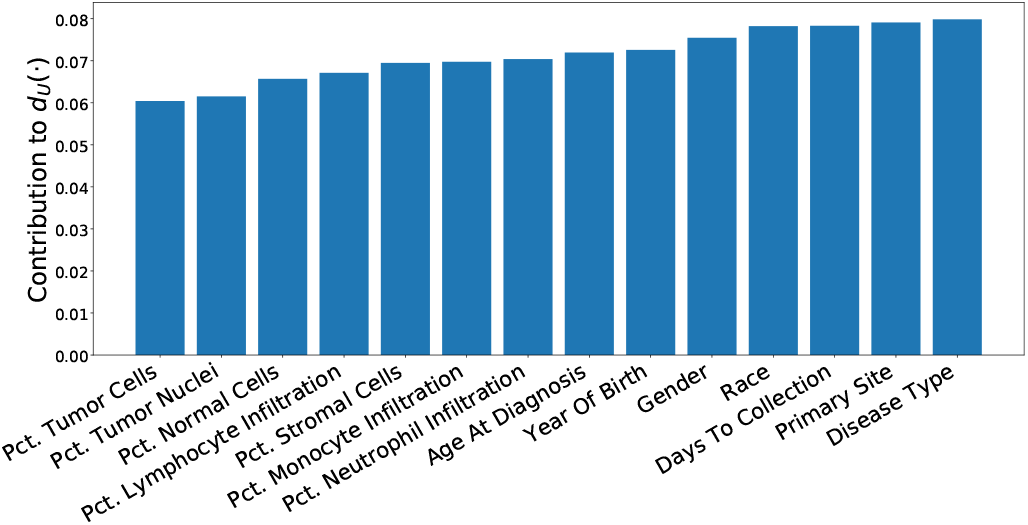
Contribution of each covariate to the learned personalization distance in the pancancer dataset. We see that, as expected, this method learns to upweight differences in disease type and primary site, along with other demographic features.

### B. Personalization Effects

We also examine the learned distance metrics for contributions to personalization by each covariate. The linear form of the distance metric makes interpretation of *ϕ*_*U*_ straightforward by inspection of the loadings (Figure 2). As expected, the disease and primary tissue site of the sample have the heaviest influence on personalization, confirming our intuition that the variation between cell types is highest in cells of distinct differentiations. Next in importance to *ϕ*_*U*_ are demographic and clinical features, which may be interpreted as a coarse-grained view of the patient’s SNPs. Important molecular markers of cancer subtype appear to be (a) percent of neutrophil infiltration, (b) percent monocyte infiltration, and (c) percent stromal cells, confirming clinical findings these phenotypic characteristics as indicative of molecular subtypes, especially in breast cancers (23–25).

### C. Accurate Recovery of Personalized Parameters

Personalized regression selects variables on a sample-specific level. Such fine-grained analytic power, unobscured by cohort averaging, enables more accurate recovery of important features than is possible by population-scale models. As a result, the number of variables selected for each sample-specific model is much lower than the number of variables selected by the population estimator (Figure 3, top). In addition, the number of samples for which each variable is selected follow a long-tailed distribution in which a few genes are selected for many samples, but many genes are selected for a few samples (Figure 3, bottom). The set of common gene selections represents well-studied oncogenes that are common to many types of cancer while the infrequently selected genes may correspond to less common oncogenes.

**Fig. 3.**
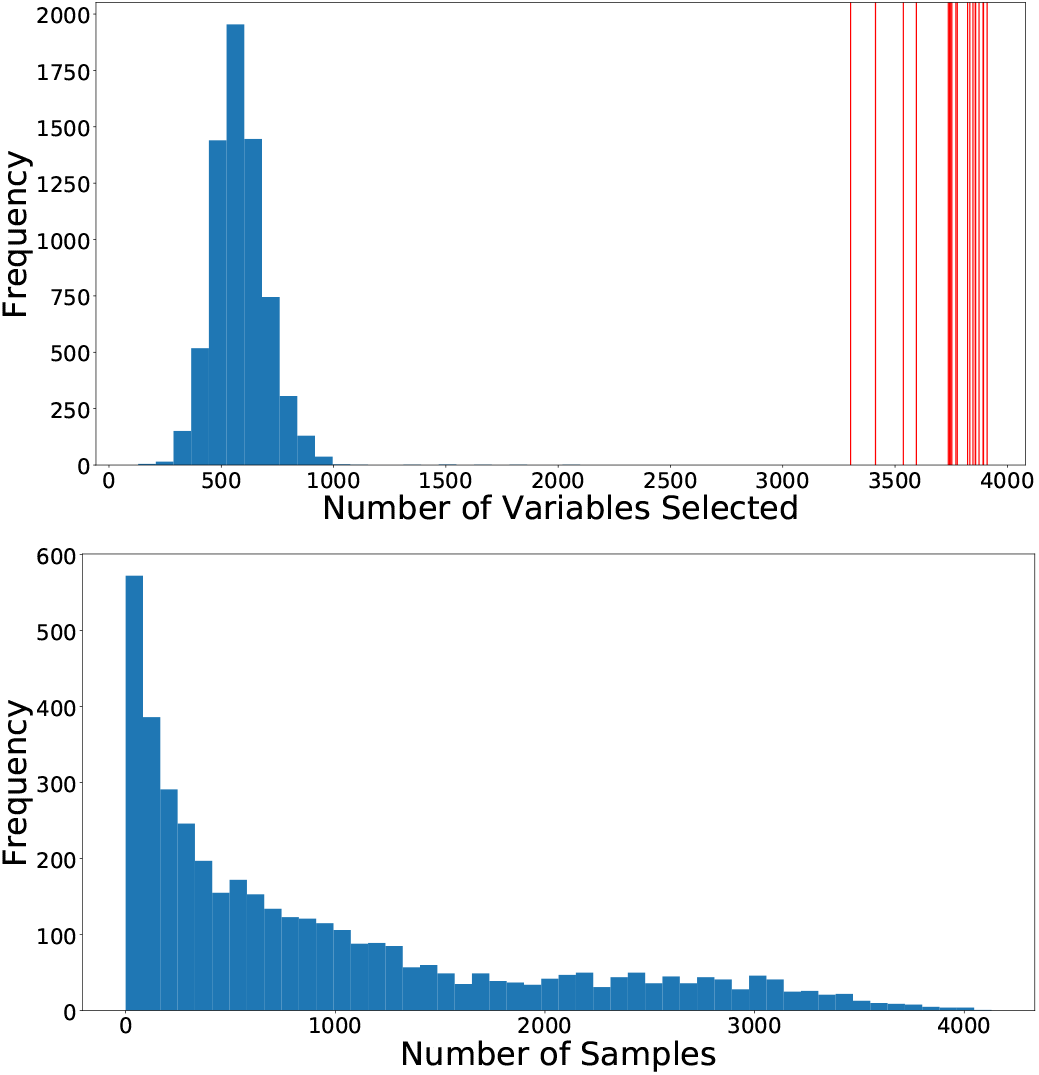
The sample-specific variable selection of personalized regression results in models with fewer selected variables than those selected by population-level models. (Top) Histogram of the number of variables selected for each patient by personalized regression. Vertical red lines indicate the number of variables selected by the Tissue-Population model trained on a single cancer type. Personalized models achieve similar or improved predictive performance with fewer selected genes. (Bottom) Histogram of the number of samples for which each gene is selected.

To investigate this possibility of many infrequently selected oncogenes, we further examine the oncogene distribution by rank of variable. Ranks are calculated by ordering the sums of the magnitudes of each coefficient along the sample axis (for population models, this is simply the magnitude of the coefficient associated with that variable). In this way, the rank captures both the number of samples for which the variable was selected and the magnitude of the implied effect size. As shown in Figure 4, the overlap between selected genetic markers and the annotations in COSMIC (21) is improved by the process of personalization. We see that the highly ranked oncogenes are efficiently selected by nearly all methods, but the performance of the baseline models lags as the rank diminishes. In particular, although the Tissue-Population models that are learned independently using only samples from a given tissue tend to select highly ranked genes that are also annotated in COSMIC, the performance in the long-tail of infrequently selected genes is less competitive compared to the personalized model. This confirms the intuition that personalization is the most useful in this latter regime.

**Fig. 4.**
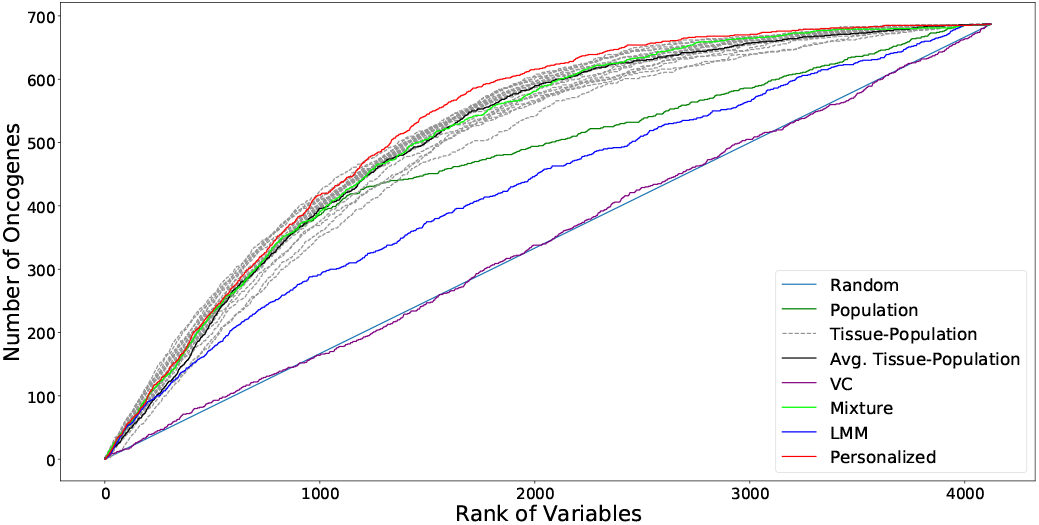
Overlap of selected variables with annotated oncogenes (best viewed in color). Results for each tissue-specific model are displayed in dashed gray lines, with the sample-weighted mean displayed in a solid black line. We see that the personalized models select oncogenes at higher ranks than do the baseline methods, especially for the long tail of low rank oncogenes.

To test whether this increase in oncogene selection is due to novel identification of genetic processes, we perform enrichment analysis of the ranked lists of genes. Reported in Table 3 are the most significant Gene Ontology (GO) terms from a ranked enrichment test using Panther 13.1 (26) on the Panther GO-SLIM Biological Process dataset (27) with a cutoff of *p* < 0.05 for the Bonferroni-corrected *p*-values. The genes selected by personalized models are enriched with similar GO terms compared to the baseline models, which is expected since the gene ontology is largely comprised of well-studied annotations from large cohorts as opposed to harder to detect personalized effects. This validates our hypothesis that the improved performance of variable selection is not due to identification of a single group of genes, but rather is due to the identification of many sample-specific effects.

**Table 3.**
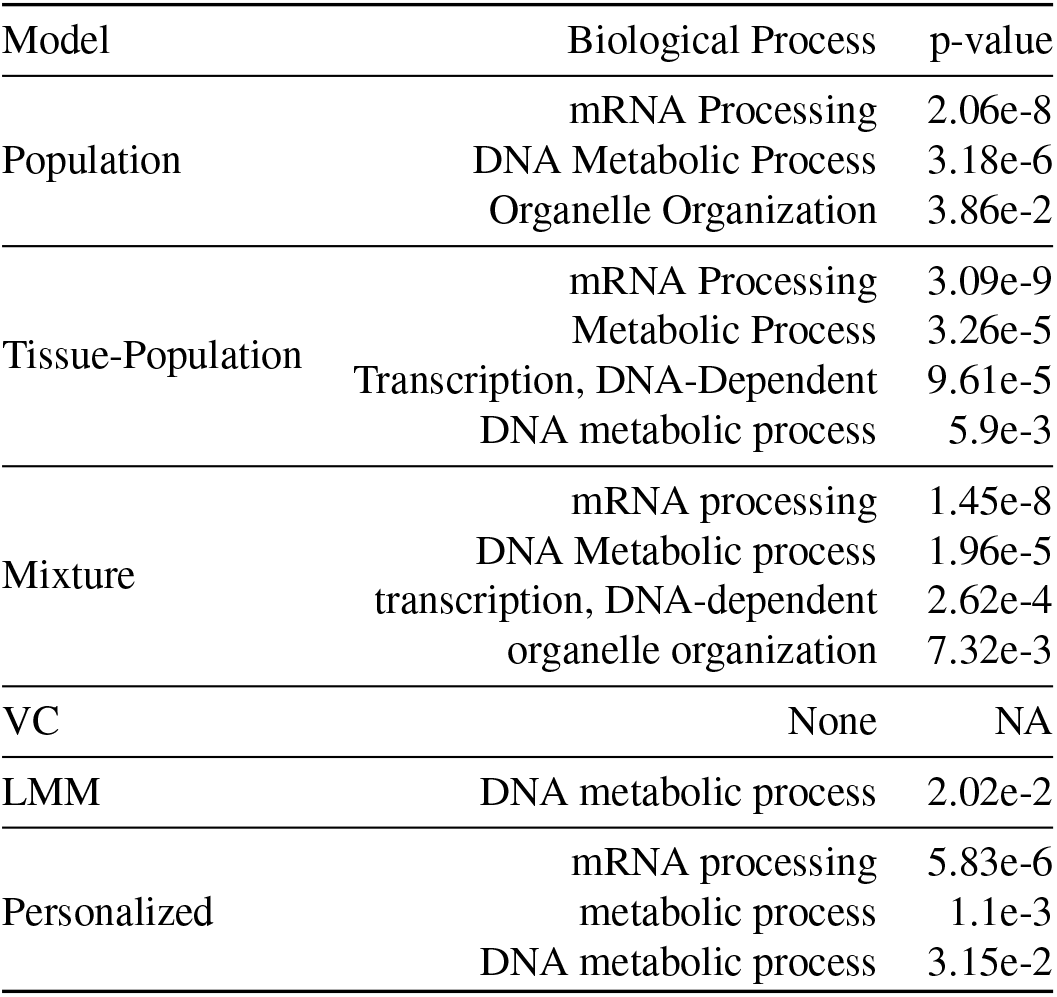
Enrichment Analysis of Complete Variable Rankings

### D. Discovery of Molecular Subtypes

The pattern of selection of genes is of particular interest for clinical application. As seen in Figure 5, there are a number of common oncogenes that are repeatedly selected throughout many cancer types, including FOXA1, HOXC13, and FCGR2B. This set combines with a sparse selection of a number of oncogenes specific to each cancer type. These cancer types span surface-level characteristics such as tissue type. Interestingly, we also see a small set of rarely selected oncogenes that are consistently selected for a cluster of about 300 patients (outlined in Figure 5). This set of oncogenes is highly over-represented for the GO biological process term “Modulation of Chemical Synaptic Transmission” (Bonferroni corrected *p*-values of 2.32e-2), which includes genes ATP1A2, SLC6A4, ASIC1, GRM3, and SLC8A3. These genes code for ion-transport processes, which have long been seen in vivo as an important system in thyroid cancer (28) and in vitro from leukemic cells (29), but only recently been appreciated as a functional marker across many different cancer types (30).

**Fig. 5.**
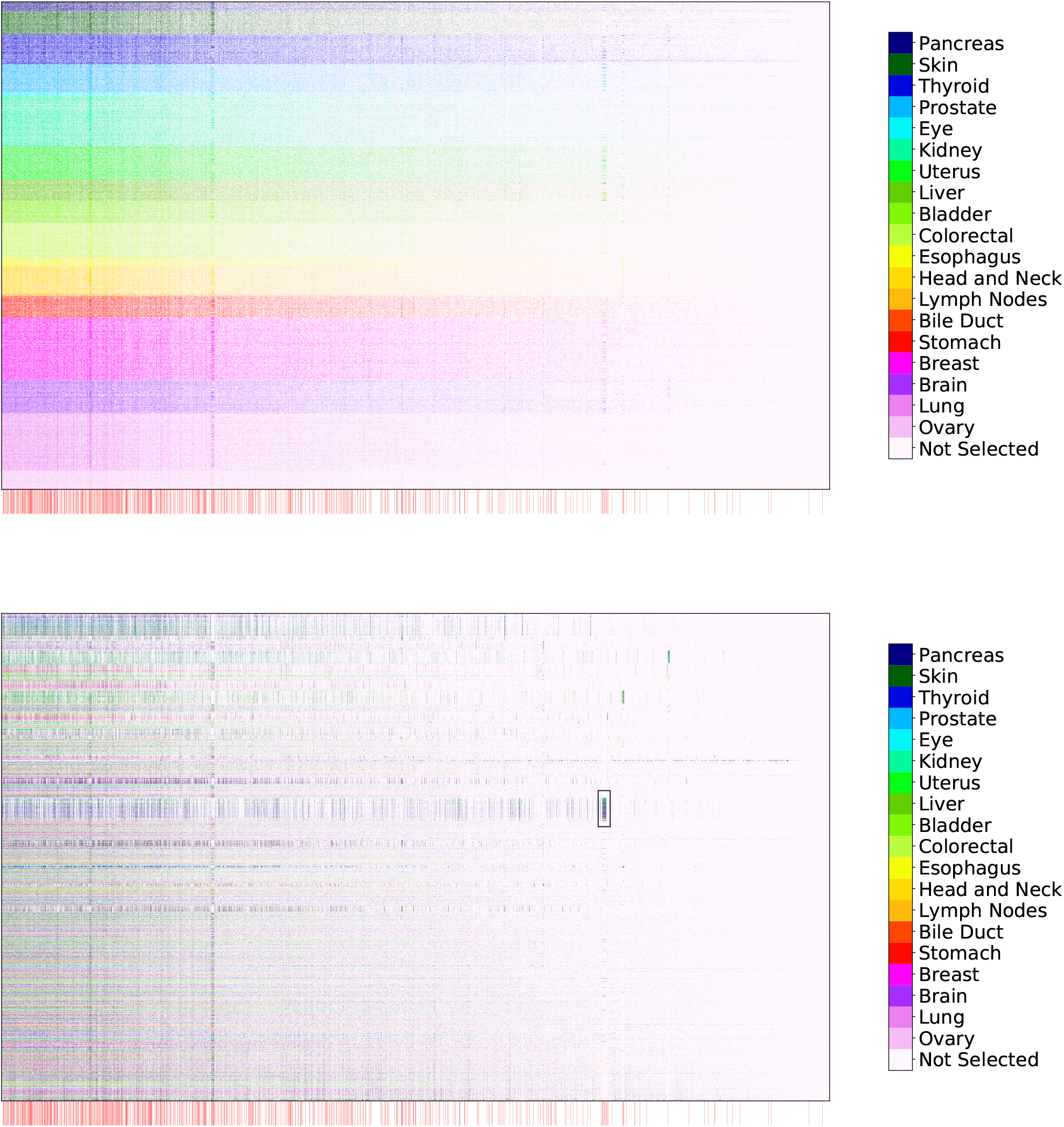
Selection of genetic markers as predictive of case/control status from a pan-cancer dataset. The horizontal axis denotes genes while the vertical axis indexes samples. Selected variables in each row are colored by the primary tumor site of the sample, with unselected variables colored white. We observe consistent selection of a number of common oncogenes throughout all cancer types along with the sparse selection of a small number of oncogenes specific to each cancer type. Genes annotated as oncogenes in the COSMIC census are marked by a red line along the horizontal axis (zoom in for more detail as these lines may be difficult to differentiate on some screens). (Top) Rows ordered by primary tissue site, (Bottom) Rows clustered according to personalized variable selection. The boxed region is analyzed in Section D.

Figure 6 depicts a tSNE projection of the learned effect vector for each sample, colored by the primary tumor site. While the samples appear to form clusters, and the case samples are separated from the control samples by a large margin, again these clusters do not appear to correspond to any individual covariate. This complexity of personalization underscores the need for learned distant metrics to capture relationships corresponding to molecular characterization of tumors.

**Fig. 6.**
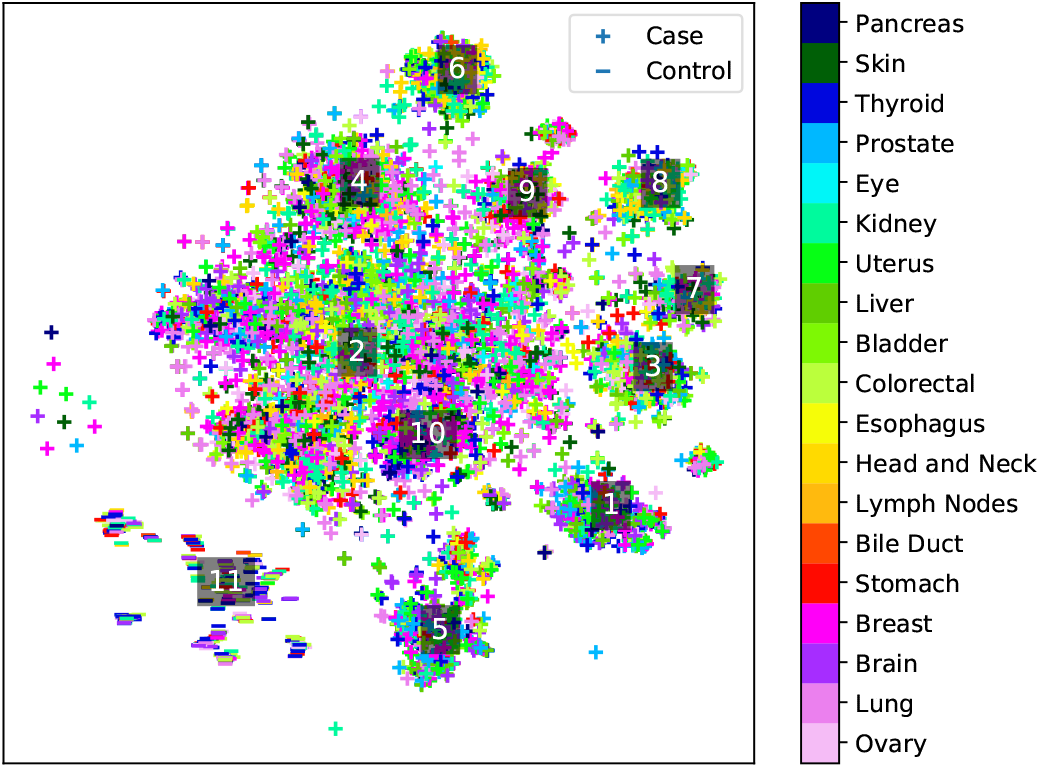
tSNE projection of personalized regression parameters learned from a pan-cancer dataset. Each point represents a single sample with color indicating primary tumor site and marker type indicating case/control status of the patient. Labelled points indicate the centroids of clusters analyzed in Table 4.

To identify molecular subtypes, we cluster the parameter embeddings using the HDBSCAN algorithm and perform an enrichment analysis of each cluster’s variable selection in an analogous manner to the procedure described in Section C. The top 3 over-enriched leaf terms from the GO biological process dataset are shown in Table 4. We see that the different clusters of models correspond to different biological processes. For instance, cluster 3 is enriched for several terms associated with extracellular interactions, while cluster 2 emphasizes terms associated with nucleotide modification via splicing and repair. These results suggest that the clusters discovered by personalized regression may correspond to clinically meaningful molecular subtypes.

**Table 4.**
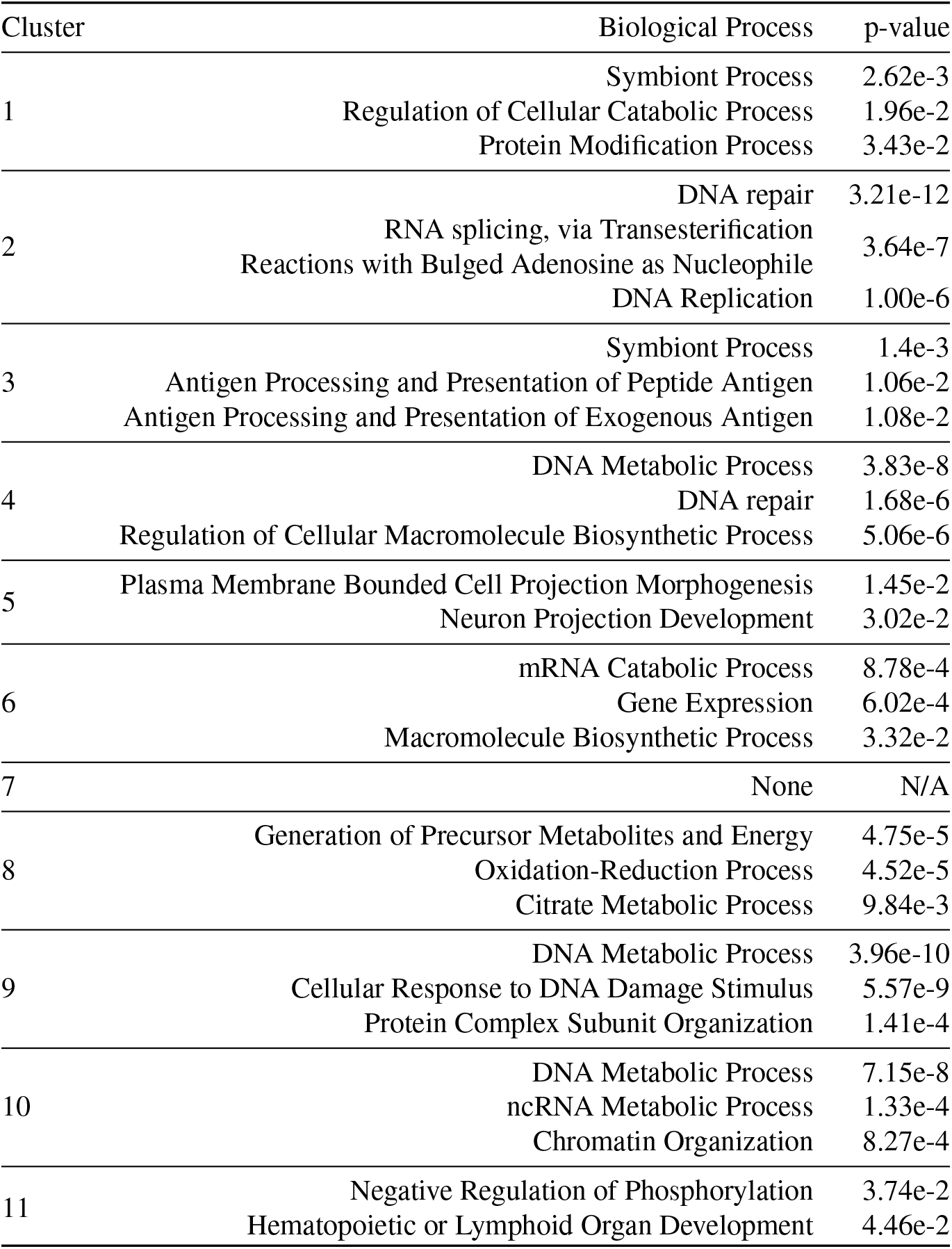
Enrichment Analysis of Tumor Clusters

## 7: Conclusions and Future Work

In this work, we have presented a framework for estimating sample-specific regression models via the introduction of a novel regularizer that matches distance in covariate values to distance in regression parameters. We have demonstrated the effectiveness of this paradigm for sample-specific tumor analysis by gene selection on a pan-cancer dataset. Much work remains to be done in the application of this method to cancer analysis. We are particularly interested in the potential to uncover novel molecular subtypes that correspond to shared mutational patterns of tumors, especially for analysis of the long tail of understudied genetic factors. In addition, we would like to apply this paradigm of sample-specific estimation to more complicated models. With the increasing number of biological assays for precise granularity buoyed by the rising tide of genomic data availability, we anticipate sample-specific modeling to continue to increase in importance and relevance to the bioinformatics community.

## Acknowledgements

We thank Maruan Al-Shedivat, Avinava Dubey, and Michael Kleyman for insightful discussion, and anonymous reviewers for constructive criticism.

## Footnotes

1 cancergenome.nih.giv

2 dcc.icgc.org

